# Slice, Subtract, Segment: Efficient 2D and Physiologically Informed 3D Brain Tumor Segmentation

**DOI:** 10.64898/2026.10.04.756528

**Authors:** Latchan Chhetri, Ayush Anand

**Author notes:** Corresponding author: Latchan Chhetri. These authors contributed equally to this work.

## Abstract

**Background:** Automated segmentation of glioma subregions on multiparametric MRI can be improved by changing either the network architecture or the information presented to the network. These two design levers are usually studied together, which makes their individual contributions difficult to see. We present an integrated analysis of two completed, independent studies, each of which isolates one lever.

**Methods:** In the architecture-level study (*Slice*), SIBA-UNet—a 2D U-Net combining depthwise separable convolution, inverted-bottleneck blocks and attention-guided skip fusion—was trained on 128*×*128 multimodal slices from BraTS 2020 with a patient-level split, and a sigmoid decision threshold was selected on validation data. In the representation-level study (*Subtract*), CMSR-3D augmented the four standard MRI sequences with four voxel-wise cross-modal subtraction maps (T1*−*T1CE, T2*−*FLAIR, FLAIR*−*T1CE, T2*−*T1CE) and evaluated 4-, 6- and 8-channel inputs with an identical 3D U-Net on 1,251 BraTS-GLI 2023 cases (1,001/125/125 split).

**Results:** SIBA-UNet reached an overall validation Dice of 0.8601 at a validation-selected threshold of 0.51, with reported subregion Dice of 0.8795 (necrotic core), 0.8501 (edema) and 0.9125 (enhancing tumor); the source ablation was only partially reported. With the architecture held fixed, the 8-channel CMSR-3D configuration gave the highest mean Dice among the evaluated inputs (0.8699 vs. 0.8534 for the 4-channel baseline) and the lowest mean HD95 (5.9859 vs. 8.2308), whereas either two-map subset alone changed mean Dice by at most 0.0037.

**Conclusions:** Architecture design and input-representation design are separable levers for brain tumor segmentation. The controlled representation ablation indicates that the complete set of subtraction maps was more informative than either subset under a fixed 3D U-Net. Both studies were evaluated internally on their respective datasets, and their numerical results are not directly compared because the datasets, dimensionality and evaluation protocols differ.

## 1 Introduction

Gliomas are the most common primary malignant tumors of the brain, and their management depends on accurate characterisation of tumor extent on magnetic resonance imaging (MRI) [1]. Routine multiparametric MRI combines complementary sequences: T1-weighted images before (T1) and after (T1CE) gadolinium contrast, which reveal enhancing tumor where the blood–brain barrier is disrupted, and T2-weighted (T2) and fluid-attenuated inversion recovery (FLAIR) images, which show the surrounding edema and non-enhancing abnormality. Delineating the necrotic core, the enhancing tumor and the peritumoral edema supports volumetric assessment and treatment planning, but manual segmentation in three dimensions is time-consuming and variable between readers [2]. The BraTS benchmark has made automated subregion segmentation a standard, openly evaluated problem [2–5], and encoder–decoder convolutional networks derived from U-Net [6, 7] remain the dominant approach [8].

Efforts to improve such networks usually pull one of two levers. The first is the *architecture*: how features are computed, how resolution is traded for context, and how encoder information reaches the decoder. Examples include attention gates on skip connections [9, 10], efficient convolutional building blocks [11, 12] and transformer encoders [13]. The second is the *input representation*: what the network is shown. In multimodal MRI, the sequences are almost always stacked as channels and the network is left to discover their relationships. Yet radiologists read glioma MRI comparatively—enhancement is judged by comparing T1CE with T1 [14], and the relationship between T2 and FLAIR signal carries diagnostic information [15]—which raises the question of whether making such comparisons explicit helps a network. In practice, published methods tend to change architecture, representation and training simultaneously, which makes the contribution of each lever hard to isolate.

This manuscript integrates two completed and independent studies, each of which isolates one lever, under a common framing that we summarise as *Slice, Subtract, Segment* (Fig. 1). In the first study (*Slice*), the input was held at the four standard sequences and the architecture of a 2D slice-based U-Net was modified. We refer to the resulting network as SIBA-UNet, a Separable Inverted-Bottleneck Attention U-Net that combines depthwise separable convolution, inverted-bottleneck blocks and attention-guided skip fusion. In the second study (*Subtract*), the architecture of a 3D U-Net was held fixed and the input representation was varied by adding voxel-wise cross-modal subtraction maps. We refer to this as CMSR-3D, a Cross-Modal Subtraction Representation evaluated in 3D. Both studies address the same task (Segment): delineating glioma subregions using the same four MRI modality types, T1, T1CE, T2 and FLAIR. Both model names are introduced here for clarity and denote the methods exactly as evaluated in the original studies.

**Figure 1.**
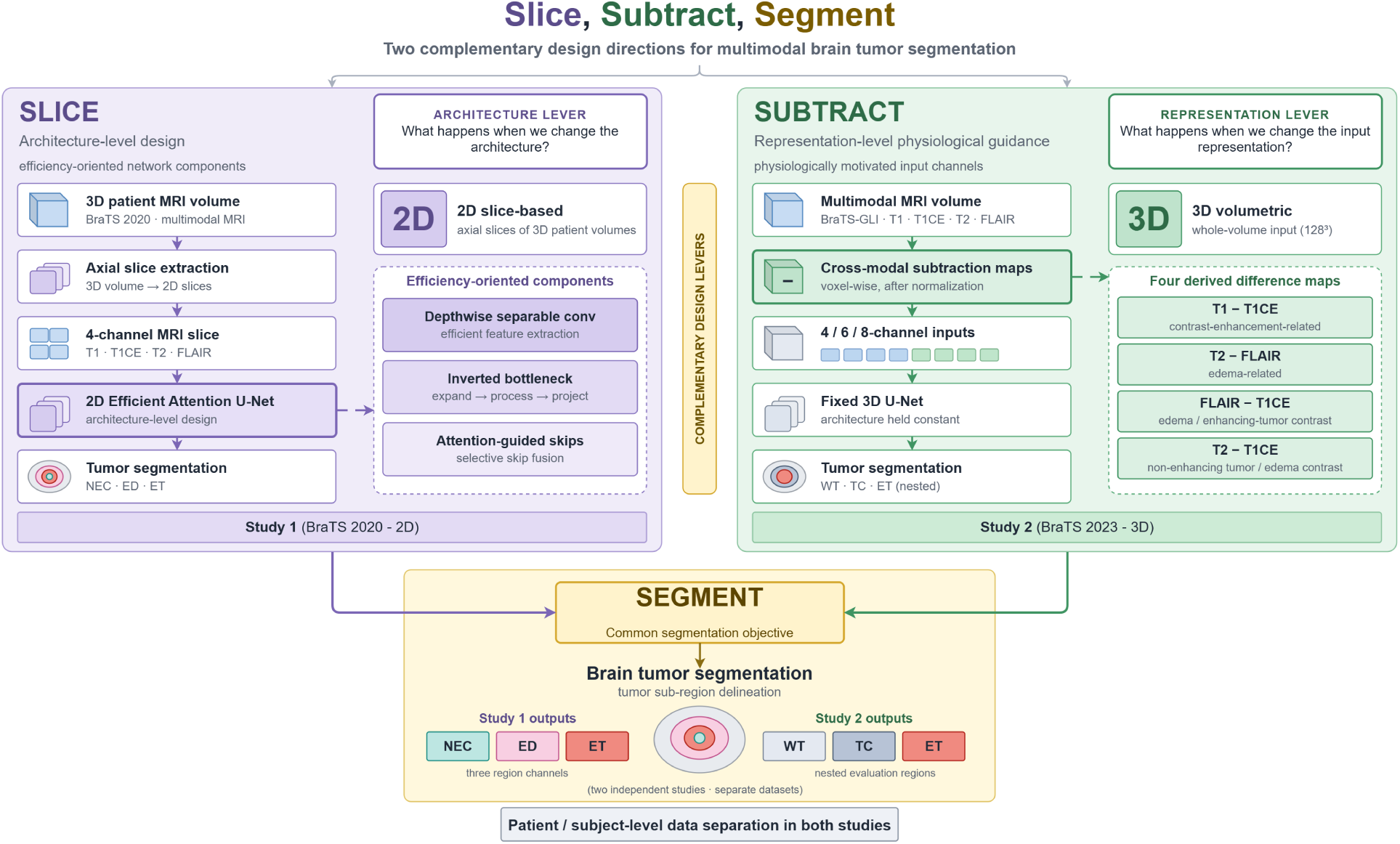
Slice, Subtract, Segment: two complementary design levers for multimodal brain tumor segmentation. Left (*Slice*, Study 1): an architecture-level design, in which 2D axial slices of the four standard MRI sequences are segmented by SIBA-UNet, whose efficiency-oriented components are depthwise separable convolution, inverted-bottleneck blocks and attention-guided skip fusion. Right (*Subtract*, Study 2): a representation-level design, in which four cross-modal subtraction maps are added to the four sequences (4-, 6- or 8-channel inputs) and evaluated with a fixed 3D U-Net (CMSR-3D). Bottom (*Segment*): the shared objective of tumor subregion delineation. Study 1 outputs three subregion channels (NEC, ED, ET); Study 2 is evaluated on the nested regions WT, TC and ET. The two studies were conducted independently on separate BraTS releases with patient- or case-level separation of training and evaluation data; the figure is conceptual and does not depict a joint model. Colours are visual encodings only. Label key: “2D Efficient Attention U-Net” in the diagram is SIBA-UNet; “Patient / subject-level data separation in both studies” refers to the patient-level split of Study 1 and the case-level split of Study 2.

The contribution of this manuscript is therefore not a new unified model but an analytical one. We (i) describe both methods in a common notation and with consistent terminology; (ii) report their results exactly as documented, stating explicitly what each source did and did not establish; (iii) contrast the two studies by the variable each manipulated and the experimental control each achieved, without comparing scores across non-comparable protocols; and (iv) draw design lessons that neither study yields alone. Among these, Study 2’s controlled ablation shows that, with the architecture fixed, the complete set of four subtraction maps was associated with the highest Dice and the lowest HD95 in every evaluation region, whereas either half of the maps alone gave at most marginal and mixed changes.

The remainder of the paper is organised as follows. Section 2 reviews related work. Section 3 describes both studies and the reporting conventions used throughout. Section 4 presents the results of each study, Section 5 analyses the two studies together, and Sections 6 to 8 discuss implications, limitations and conclusions.

## 2 Related Work

### Brain tumor segmentation benchmarks

The BraTS challenge established a common multiparametric MRI dataset and evaluation protocol for glioma subregion segmentation [2]. It was subsequently extended with expert annotations of The Cancer Genome Atlas collections [3], with multi-year evaluations of segmentation and related tasks [4], and with the substantially larger BraTS 2021 collection [5], on which the 2023 adult glioma data build. Across releases, performance is reported on nested evaluation regions (whole tumor, tumor core and enhancing tumor) using Dice and the 95th-percentile Hausdorff distance [2, 5]. Challenge rankings depend strongly on the evaluation protocol and metric aggregation [16]. This is one reason why results obtained on internal splits and on official test sets, or on different releases, are not directly comparable.

### Encoder-decoder architectures

U-Net introduced the symmetric encoder–decoder with skip connections that underlies most biomedical segmentation networks [6]. It was extended to volumetric data as 3D U-Net [7] and, with residual blocks and a Dice-based objective, as V-Net [17]; nested skip pathways were explored in UNet++ [18]. For brain tumors specifically, early deep models used multi-path 2D networks [19] and multi-scale 3D networks with conditional random fields [20], and later high-performing BraTS entries included an autoencoder-regularised 3D network (the BraTS 2018 winner) [21] and transformer encoders such as Swin UNETR [13]. The self-configuring nnU-Net showed that a carefully configured standard U-Net remains highly competitive across tasks [8]. This underlines the importance of controlling the training pipeline when attributing improvements to architectural changes. Deep supervision of intermediate decoder outputs [22] and instance normalisation [23] are common components of such pipelines and are used in Study 2.

### Attention gating and efficient building blocks

Attention U-Net introduced additive attention gates that reweight skip features using a gating signal from the coarser decoder level, suppressing irrelevant regions without external localisation [9, 10]. Separately, depthwise separable convolutions factorise spatial and channel mixing to reduce computation [11, 24]. MobileNetV2 combined them with inverted residual blocks that expand to a wide intermediate representation and project linearly back to a narrow one [12]. Efficiency-oriented segmentation networks such as UNeXt adapt such ideas to medical images [25]. Study 1 combines separable inverted-bottleneck blocks with attention-gated additive skip fusion in a 2D slice-based U-Net.

### 2D versus 3D formulations

Slice-based 2D models are memory-efficient and can use large batches, but they lack through-plane context; 3D models capture volumetric context at greater memory cost and usually require cropping or patching. A direct comparison on brain MRI segmentation found that 3D models were generally more accurate, while 2D and 2.5D models required less memory [26]. The two studies integrated here sit on either side of this trade-off.

### Input representation and cross-modal contrast

Most deep segmentation methods present the MRI sequences as stacked channels and leave cross-sequence relationships to be learned. In radiological practice, however, explicit comparisons between sequences are informative. T1-weighted subtraction maps (post-minus pre-contrast) improve the delineation of enhancing tumor [14], and the relationship between T2 and FLAIR signal (the T2–FLAIR mismatch sign) is associated with molecular glioma subtype [15]. These observations motivate Study 2, which supplies voxel-wise differences between normalised sequences as additional input channels and evaluates their contribution with the architecture held fixed.

### Evaluation and reporting

Recent consensus work has catalogued metric-related pitfalls in image analysis validation and recommended matching metrics to the clinical question [27, 28]. Methodological reviews have highlighted the effect of unreported variance and of validation–test conflation on the conclusions of medical imaging studies [29]. The Checklist for Artificial Intelligence in Medical Imaging (CLAIM) provides reporting guidance for such studies [30, 31]. These works inform the reporting conventions of the present manuscript.

## 3 Materials and Methods

### 3.1 Overall study design and data sources

This manuscript integrates two completed and independently conducted studies of multimodal brain tumor segmentation. Each study varied one design lever while leaving the other unchanged in kind. Study 1 (*Slice*) modified the network architecture of a 2D slice-based U-Net while keeping the four standard MRI sequences as input. Study 2 (*Subtract*) modified the input representation of a 3D volumetric U-Net while holding the architecture fixed. The two models were not trained jointly, did not share data splits, and were evaluated under different label conventions; no experiment combining them was performed for this manuscript. All methodological details below are taken from the original study reports; where a detail was not documented, this is stated explicitly. Table 1 summarises both studies side by side.

**Table 1.** Design of the two integrated studies. Entries are taken from the original study reports.

|  | **Study 1 — *Slice*** | **Study 2 — *Subtract*** |
| --- | --- | --- |
| Design lever varied | Network architecture | Input representation |
| Held constant | Four standard MRI inputs | 3D U-Net architecture, split, training protocol |
| Model / framework | SIBA-UNet (2D) | CMSR-3D with fixed 3D U-Net |
| Dataset | BraTS 2020 (training collection) | BraTS-GLI 2023; 1,251 cases |
| Dimensionality | 2D axial slices, $128 \times 128$; 75 slices per volume from index 40 | 3D volumes, $128 \times 128 \times 128$ (centre crop / zero pad) |
| Input channels | 4 (T1, T1CE, T2, FLAIR) | 4, 6 (Set A or B) or 8 (4 modalities + 4 difference maps) |
| Intensity normalisation | Min–max, per sample | z-score per modality, non-zero voxels |
| Data partition | Patient-level, 70:15:15; | Case-level, 1,001 / 125 / 125 (80:10:10), seed 42 |
| Augmentation | None | None |
| Model output | 3 sigmoid channels: NEC, ED, ET (multi-label) | 4 softmax classes: BG, NCR, ED, ET |
| Evaluation targets | NEC, ED, ET; “overall” | Nested regions WT, TC, ET; means |
| Loss | - | Foreground soft-Dice + cross-entropy (equal weights); deep supervision 0.5 / 0.25 |
| Optimiser and schedule | Adam, $10^{-4}$, $\times 0.98$ per epoch; 200 epochs; batch 32 | AdamW, $10^{-4}$, weight decay $10^{-5}$; cosine to $10^{-6}$; 200 epochs; batch 1 |
| Model selection | Threshold 0.51 selected on validation Dice | Checkpoint with best validation mean Dice |
| Metrics | Dice | Dice, HD95 |
| Reported partition | Validation | Held-out (validation or test not specified) |
| Runs per configuration | 1 | 1 |

Both studies used publicly released, de-identified data from the Brain Tumor Segmentation (BraTS) benchmark [2–5]. Study 1 used the BraTS 2020 training collection; Study 2 used the BraTS 2023 adult glioma (BraTS-GLI) collection, which builds on the BraTS 2021 data [5]. The two studies use different BraTS releases and should not be treated as independent cohorts for cross-study numerical comparison. Each case comprises co-registered, skull-stripped T1-weighted (T1), post-contrast T1-weighted (T1CE), T2-weighted (T2) and T2 fluid-attenuated inversion recovery (FLAIR) volumes with expert annotations of the tumor subregions [2, 5].

### 3.2 Study 1: Efficient 2D segmentation

#### 3.2.1 Dataset and slice preparation

Study 1 processed each BraTS 2020 volume as a stack of 2D axial slices. For every volume, 75 consecutive axial slices beginning at slice index 40 were retained; this fixed window concentrates training on the central portion of the brain, where tumor tissue most often lies, and reduces computational cost. The window was fixed rather than selected per case, so it does not guarantee that every retained slice contains tumor. Each retained slice was resized to 128 *×* 128 pixels. The four co-registered modalities were stacked as channels, giving an input tensor of shape *B×* 4 *×* 128 *×* 128 for a batch of size *B*. Segmentation masks were resized with nearest-neighbour interpolation to preserve label values and converted to a three-channel multi-label target encoding the necrotic core (NEC), edema (ED) and enhancing tumor (ET). Intensities were scaled per sample with min–max normalisation. Data were divided at the patient level into training, validation and test partitions in an approximately 70:15:15 ratio, so that no patient contributed slices to more than one partition; the exact number of patients per partition was not reported in the source. Data augmentation was not used.

#### 3.2.2 SIBA-UNet architecture

We refer to the Study 1 network as SIBA-UNet (**S**eparable **I**nverted-**B**ottleneck **A**ttention U-Net). The name is introduced in this manuscript for clarity only; SIBA-UNet is exactly the architecture described and evaluated in the original Study 1 report, where it was referred to by several working names. Its ablation variant without attention gates is referred to as SIB-UNet.

As shown in Fig. 2, SIBA-UNet follows the encoder–bottleneck–decoder layout of U-Net [6] with four resolution levels. Two modifications distinguish it from a standard 2D U-Net: (i) the encoder and decoder blocks are built from depthwise separable convolutions arranged as inverted bottlenecks [11, 12, 24], and (ii) every skip connection passes through an additive attention gate before fusion with the decoder stream [9, 10]. The network ends in a three-channel output layer with sigmoid activation, producing one probability map per subregion (NEC, ED, ET). The channel widths *C*_1_*, …, C*_4_, the bottleneck width *C_b_*, the expansion factor *e*, the kernel sizes and the downsampling and upsampling operators were not reported in the source and are therefore kept symbolic.

**Figure 2.**
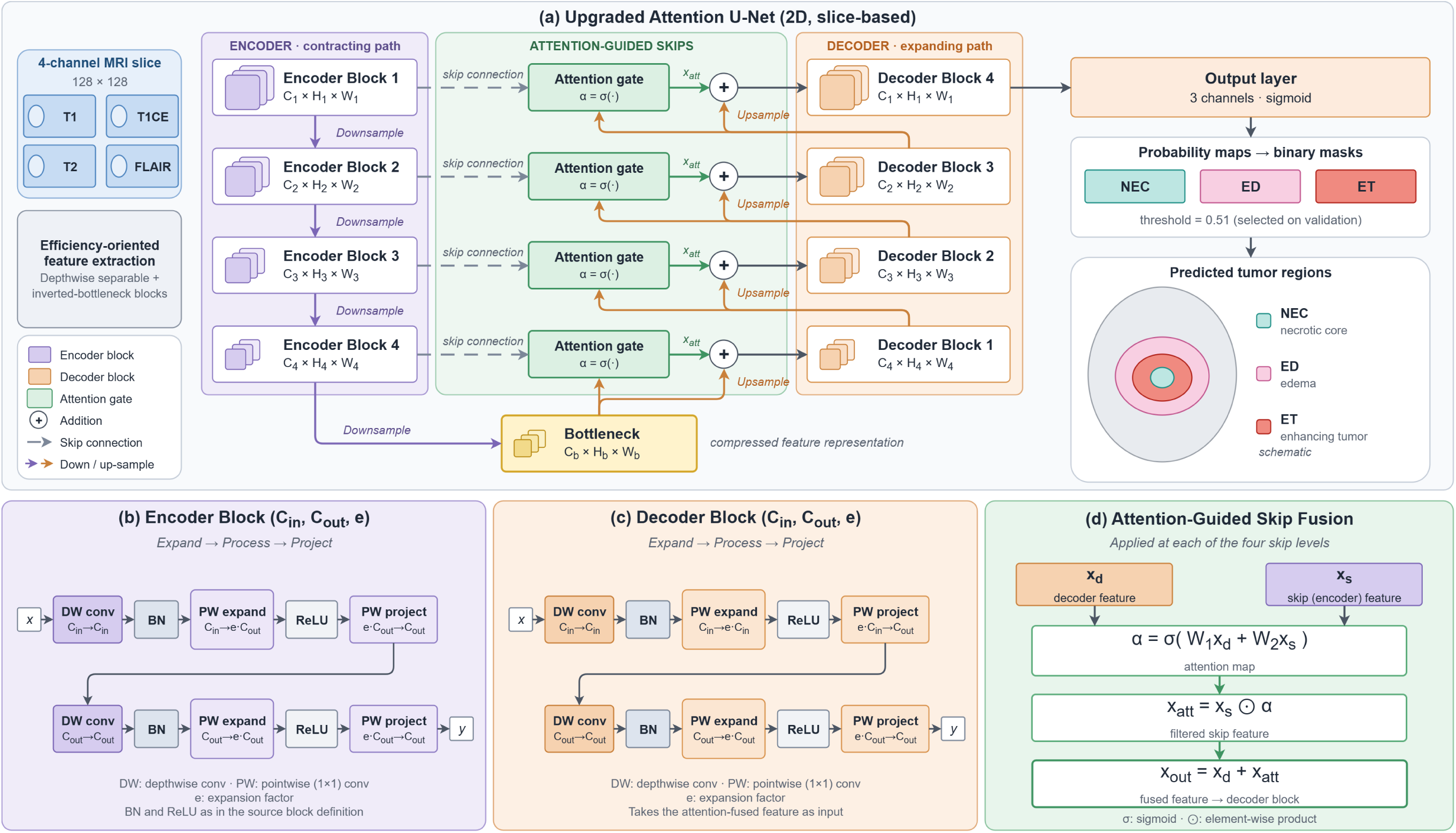
Study 1 architecture: SIBA-UNet. (a) Overall 2D slice-based encoder–decoder. A four-channel 128 *×* 128 slice (T1, T1CE, T2, FLAIR) passes through four encoder blocks and a bottleneck; each skip feature is filtered by an attention gate and added to the upsampled decoder feature before the corresponding decoder block. A three-channel sigmoid output yields probability maps that are binarised at the validation-selected threshold of 0.51. (b) Encoder block (Eq. (3)): depthwise convolution, batch normalisation, pointwise expansion, ReLU and linear pointwise projection, applied twice. (c) Decoder block (Eq. (4)), which expands relative to its input width. (d) Attention-guided skip fusion (Eqs. (5) to (7)). Channel and spatial dimensions are shown symbolically because their numerical values were not reported. In panel (a), the colours of the output boxes and of the nested schematic are visual encodings only and do not correspond to the mask colours used in Fig. 5. Label key: the model labelled “Upgraded Attention U-Net” in the diagram is SIBA-UNet; the variant without attention gates is SIB-UNet.

#### 3.2.3 Depthwise separable convolution and inverted bottleneck

For an input **x** *∈* R*^C×H×W^*, a depthwise convolution (DW) filters each channel independently with its own

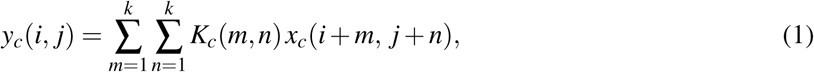

and a pointwise (1 *×* 1) convolution (PW) mixes channels,

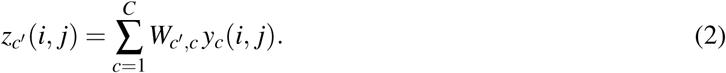

Factorising a standard convolution in this way reduces its parameter count and arithmetic cost for a given receptive field [11, 24]. Let *N* (*·*) denote batch normalisation and *φ* (*·*) the ReLU non-linearity. With expansion factor *e* and *C_e_* = *eC*_out_, an encoder block maps an input **x** with *C*_in_ channels to an output **y** with *C*_out_ channels as

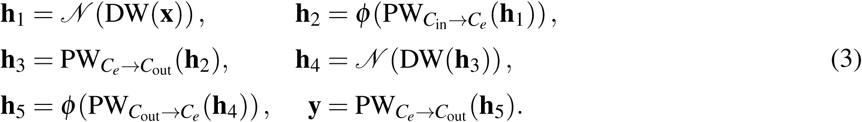

Each half of the block therefore expands the representation to a wider intermediate space, applies a non-linearity there, and projects linearly back to a narrow output, which is the inverted-bottleneck pattern of MobileNetV2 [12]. The decoder block has the same structure but expands relative to its input width, with 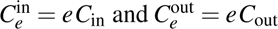:

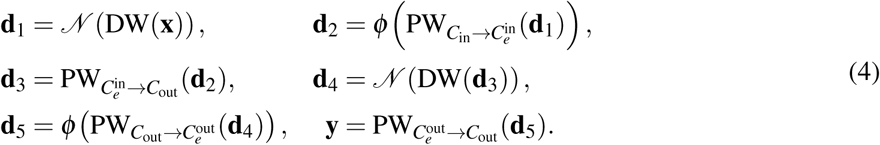

The blocks do not contain residual (identity) connections in the source description. Because *e* and the channel widths were not reported, the parameter count and computational cost of SIBA-UNet cannot be stated; we therefore describe the design as *efficiency-oriented* rather than as quantitatively more efficient.

#### 3.2.4 Attention-guided skip fusion

At each of the four skip levels, the encoder (skip) feature **x***_s_* is reweighted by an attention map computed jointly from **x***_s_* and the upsampled decoder feature **x***_d_* [9, 10]:

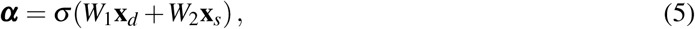

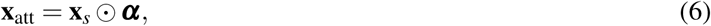

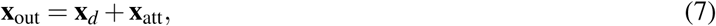

where *W*_1_ and *W*_2_ are learnable linear projections, *σ* is the logistic sigmoid and *⊙* denotes element-wise multiplication. The fused feature **x**_out_ is the input to the corresponding decoder block. Two features of this formulation differ from the original Attention U-Net gate [9]: the gating signal and skip feature are combined without an intermediate ReLU and 1 *×* 1 projection to a scalar map, and fusion is additive rather than by concatenation. Whether ***α*** is a single-channel or a channel-wise map was not specified in the source.

#### 3.2.5 Training and threshold selection

Study 1 was implemented in PyTorch. Training used the Adam optimiser [32] with an initial learning rate of 1 *×* 10^−4^, decayed exponentially by a factor of 0.98 per epoch, for 200 epochs with a batch size of 32 slices drawn from different patient volumes. The loss function used in Study 1 was not specified in the source report. Sigmoid probabilities were converted to binary masks with a single global threshold. Instead of fixing the threshold at 0.5, the source evaluated thresholds from 0.30 to 0.57 in steps of 0.03 on the validation partition and selected the value giving the highest overall validation Dice. The selected threshold of 0.51 was used for all reported Study 1 results.

### 3.3 Study 2: Physiologically informed 3D segmentation

#### 3.3.1 Dataset and preprocessing

Study 2 used 1,251 BraTS-GLI 2023 cases. Case identifiers were sorted, shuffled with a fixed random seed of 42 and split 80:10:10 into 1,001 training, 125 validation and 125 test cases. The same deterministic split was used for every input configuration, and the test partition was reserved for final evaluation. The source describes this split as subject-level. Because the identifiers in this collection carry time-point suffixes and the source does not state whether multiple time-points from the same patient were grouped before splitting, we report the documented procedure as a case-level partition and note this uncertainty as a limitation.

Each modality was z-score normalised independently using non-zero (brain) voxels only, with zero-valued background preserved. The normalised volumes were centre-cropped or zero-padded to 128 *×* 128 *×* 128 voxels. No data augmentation was used.

#### 3.3.2 Cross-modal subtraction representation

We refer to the Study 2 approach as CMSR-3D (**C**ross-**M**odal **S**ubtraction **R**epresentation, 3D). CMSR-3D denotes the input representation together with the fixed 3D U-Net used to evaluate it; it is the method described in the original Study 2 report under a different working name, and it does not introduce a new network architecture.

Let 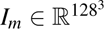 denote the normalised volume of modality *m*. Four difference maps were computed voxel-wise after normalisation:

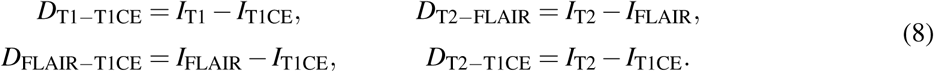

No further normalisation was applied to the difference maps. Each map was motivated by a contrast relationship used in neuro-oncological image reading. T1*−*T1CE relates to contrast enhancement, the basis of T1 subtraction imaging for delineating enhancing tumor [14]. T2*−*FLAIR relates to the signal difference produced by fluid suppression, whose spatial pattern is used qualitatively in glioma assessment [15]. FLAIR*−*T1CE contrasts FLAIR hyperintensity with enhancement, and T2*−*T1CE contrasts T2 hyperintensity with enhancement, which may help separate non-enhancing abnormality and edema from enhancing tissue. These motivations explain why the maps were chosen. They are not validated physiological measurements: after per-modality z-scoring, the maps are differences of standardised intensities rather than quantitative tissue parameters.

#### 3.3.3 Fixed 3D U-Net

All input configurations were evaluated with the same 3D U-Net [7] (Fig. 3). The encoder had four stages with 16, 32, 64 and 128 feature channels, followed by a 256-channel bottleneck at 1*/*16 of the input resolution; the decoder had four stages with transposed-convolution upsampling and skip connections. Each stage comprised two blocks of 3 *×* 3 *×* 3 convolution, instance normalisation [23] and LeakyReLU activation. A final convolution produced four output classes with softmax normalisation. Auxiliary deep-supervision heads [22] were attached to two coarser decoder outputs during training only. The downsampling operator and the exact decoder levels carrying the auxiliary heads were not specified in the source; Fig. 3 depicts them schematically. Only the number of input channels in the first convolution changed between configurations.

**Figure 3.**
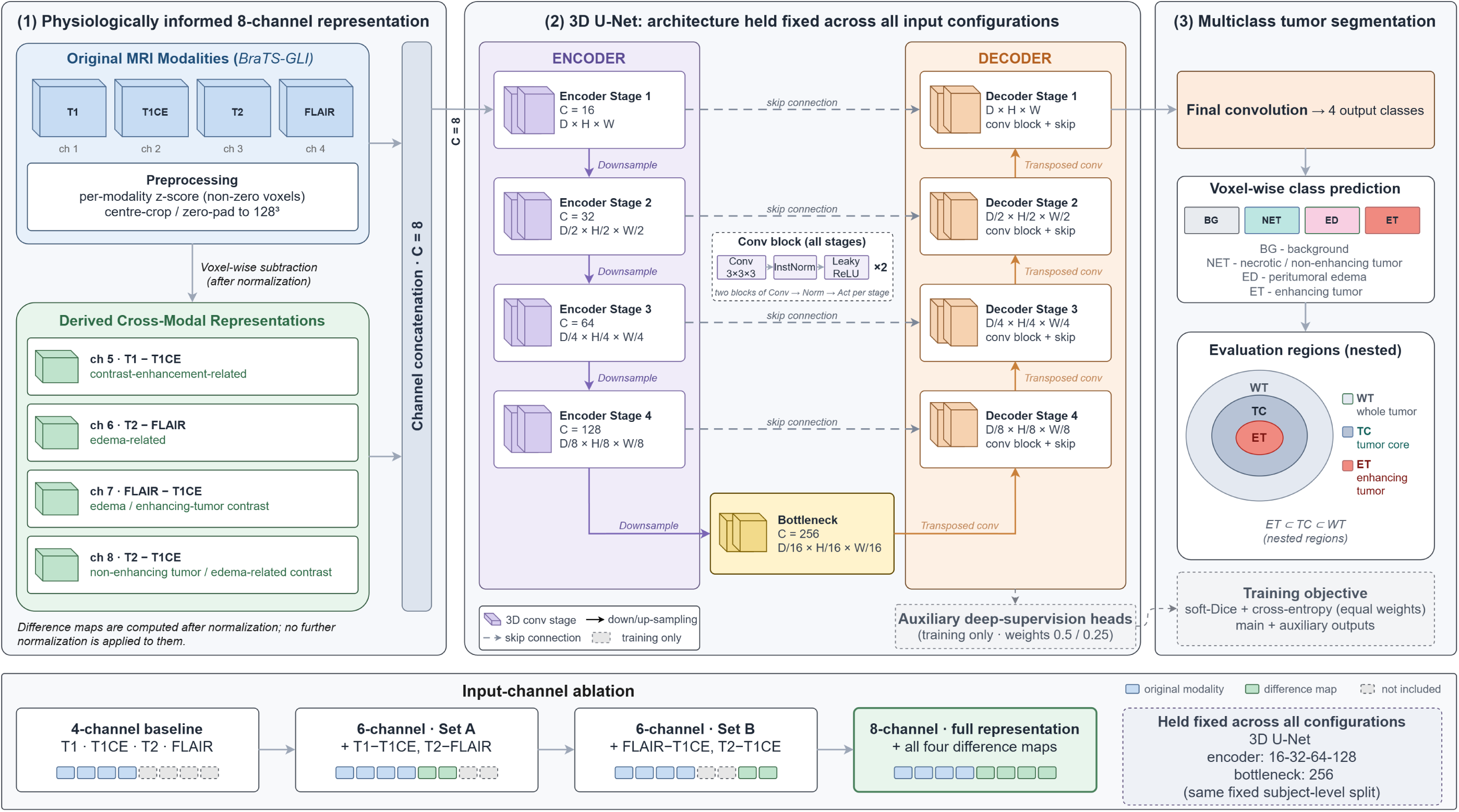
Study 2 framework: CMSR-3D with a fixed 3D U-Net. (1) The four original modalities are z-score normalised (non-zero voxels) and cropped or padded to 128^3^; four voxel-wise difference maps are then computed (Eq. (8)) and concatenated with them to form up to eight input channels. (2) The 3D U-Net (encoder widths 16–32–64–128, 256-channel bottleneck, two 3 *×* 3 *×* 3 convolution–InstanceNorm–LeakyReLU blocks per stage, transposed-convolution upsampling) is identical for all input configurations; auxiliary deep-supervision heads are used during training only. (3) The network outputs four voxel-wise classes (BG, NCR, ED, ET), from which the nested evaluation regions WT, TC and ET are derived. Bottom: the four input configurations compared in the ablation. These are independent training runs with the same split, not a sequential procedure. Colours are visual encodings only. Label key: “NET” in the diagram denotes the necrotic tumor core class, written NCR in the text; “subject-level split” refers to the case-level split described in Section 3.3.1.

The model output and the evaluation regions are distinct. The network predicts four mutually exclusive classes per voxel: background (BG), necrotic tumor core (NCR), peritumoral edema (ED) and enhancing tumor (ET), corresponding to BraTS 2023 labels 0, 1, 2 and 3. Performance was evaluated on three nested regions derived from these classes: whole tumor (WT = NCR *∪* ED *∪* ET), tumor core (TC = NCR *∪* ET) and enhancing tumor (ET), so that ET *⊂* TC *⊂* WT.

#### 3.3.4 Input-channel ablation

Four input configurations were compared (Fig. 3, bottom):

- **4-channel baseline:** T1, T1CE, T2, FLAIR;
- **6-channel Set A:** baseline + T1*−*T1CE + T2*−*FLAIR;
- **6-channel Set B:** baseline + FLAIR*−*T1CE + T2*−*T1CE;
- **8-channel (full CMSR-3D):** baseline + all four difference maps.

Set A pairs the enhancement-related and fluid-suppression-related maps; Set B pairs the two maps referenced to T1CE. Because the architecture, preprocessing, split, optimiser and checkpoint rule were identical across configurations, differences between configurations are attributable to the input representation within the limits of single training runs.

#### 3.3.5 Training procedure

Training used a batch size of 1, the AdamW optimiser [33] with an initial learning rate of 10^−4^ and weight decay of 10^−5^, cosine-annealing of the learning rate to 10^−6^[34], mixed-precision arithmetic and gradient-norm clipping at 1. The objective was the equally weighted sum of a foreground soft-Dice loss and the cross-entropy loss,

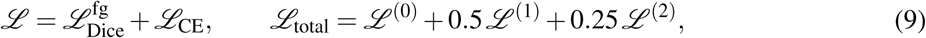

where 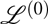 is computed on the main output and 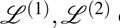 on the two auxiliary outputs. The checkpoint with the highest validation mean Dice across WT, TC and ET was retained. The model was trained for 200 epochs. Experiments were run on a single NVIDIA GeForce RTX 3050 GPU with 6 GB of memory.

### 3.4 Evaluation metrics

Segmentation overlap was measured with the Dice similarity coefficient [35],

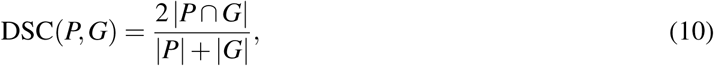

where *P* and *G* are the predicted and reference voxel (or pixel) sets for a region. Study 2 additionally reported the 95th-percentile Hausdorff distance (HD95) [36], the 95th percentile of the symmetric surface-to-surface distances between *P* and *G*, which is more sensitive to boundary errors and outlying false positives than Dice [27, 28]. Higher Dice and lower HD95 indicate better agreement.

In Study 1, Dice was computed separately for NEC, ED and ET and for the tumor as a whole. The source does not document whether Dice was aggregated per slice, per volume or over all pixels of the validation partition, nor how slices without tumor were handled. In Study 2, Dice and HD95 were reported per evaluation region (WT, TC, ET), together with their means across the three regions. The source does not specify the HD95 unit (voxels in the 128^3^ grid or millimetres), how empty predictions were scored, or whether metrics were computed in the cropped 128^3^ space or after restoring the original image geometry. HD95 values are therefore reported without units.

### 3.5 Reporting conventions

The following conventions are used throughout the manuscript.

1. **Source fidelity.** All numerical results are reproduced exactly as reported in the original study reports. No values were re-estimated from plots, imputed or recomputed, except for differences between configurations, which are simple arithmetic on reported values. Missing values are marked NR (not reported).
2. **Evaluation partitions.** Study 1 results are labelled *validation* wherever the source identifies them as such; the source states that its results were obtained on a custom validation split rather than the official BraTS test set. Study 2 ablation results are described as *held-out evaluation results*, because the source documents a reserved test partition but does not state unambiguously which partition the reported table was computed on.
3. **Single runs.** Each configuration in both studies corresponds to a single training run. No repeated-run variability, confidence intervals or significance tests were reported, and none are presented here.
4. **Label names.** Study 1 uses NEC, ED and ET for its three sigmoid output channels, following the source (BraTS 2020 label 1, the necrotic and non-enhancing tumor core, is denoted NEC). Study 2 uses BG, NCR, ED and ET for its four softmax classes, following the BraTS 2023 convention, and WT, TC and ET for the nested evaluation regions. Legends in some source-derived panels read “NCR/NET”; this denotes the same class 1 as NCR.
5. **No cross-study comparison of scores.** Because the studies differ in dataset release, dimensionality, preprocessing, output formulation, label definitions, partitions and aggregation, numerical results are never compared across studies (Section 5.2).

## 4 Results

### 4.1 Study 1: Efficient 2D segmentation

#### 4.1.1 Training behaviour and threshold selection

Fig. 4(a) shows the training and validation loss of SIBA-UNet over 200 epochs, and Fig. 4(b) shows overall validation Dice across sigmoid thresholds. Both losses fell steeply during roughly the first 50 epochs and then decreased gradually. Validation loss was markedly unstable early in training, with several spikes during the first *∼*20 epochs, and became smooth thereafter. In the second half of training the two curves ran close together, with training loss fluctuating around a slowly decreasing validation loss, and they did not diverge in a sustained way by epoch 200. Within this single train–validation split, the curves therefore give no indication of progressive overfitting. Because the loss function was not reported, the absolute loss values cannot be interpreted.

**Figure 4.**
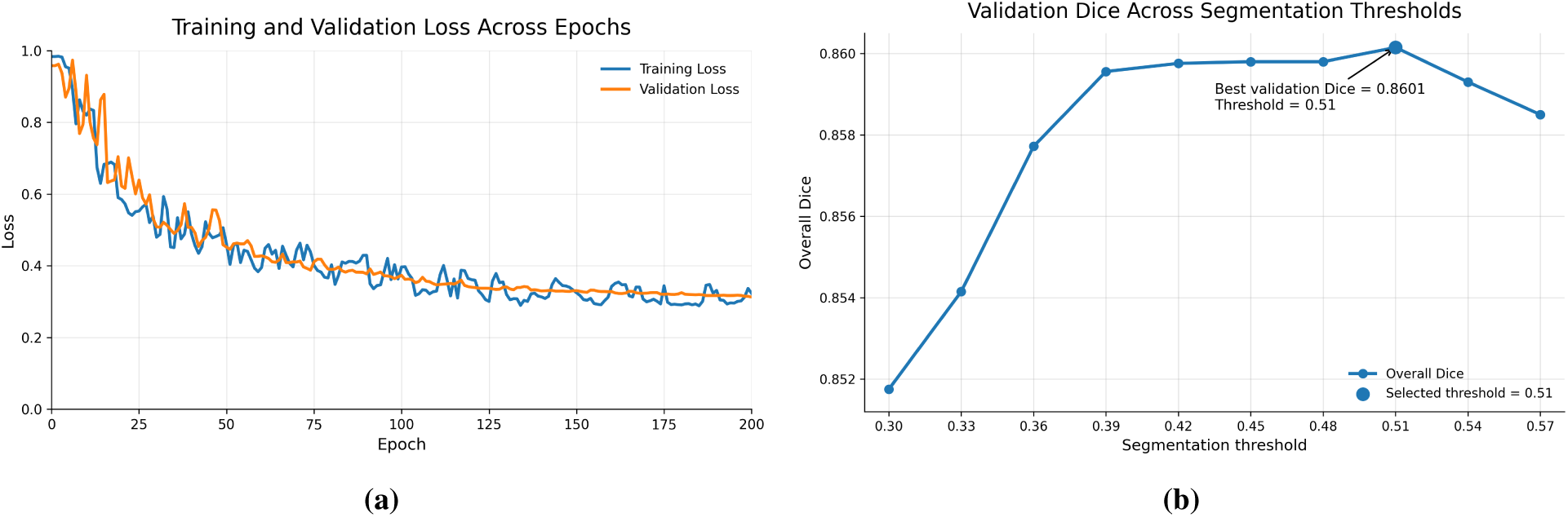
Study 1 training behaviour and threshold selection for SIBA-UNet (BraTS 2020). (a) Training and validation loss per epoch over 200 epochs, redrawn from the original study’s training history (loss function not reported). (b) Overall validation Dice as a function of the sigmoid decision threshold (0.30–0.57, step 0.03). The selected threshold of 0.51 gives the reported overall validation Dice of 0.8601. Only the selected point (threshold 0.51, Dice 0.8601) is reported numerically in the source; all other plotted values are graphical.

The threshold sweep (Fig. 4b) showed a broad plateau. Overall validation Dice rose from about 0.852 at a threshold of 0.30 to about 0.860 at 0.39, stayed within roughly 0.0007 of its maximum between 0.39 and 0.51, and declined slightly at 0.54 and 0.57. The maximum, an overall Dice of 0.8601 as reported in the source study, occurred at 0.51, which was adopted as the operating threshold. Of the plotted values, only this selected operating point is reported numerically in the source. Because the threshold and the reported Dice were selected and measured on the same validation partition, 0.8601 is an optimistically selected validation estimate rather than a held-out test result. The plotted curve suggests a relatively broad plateau around the selected threshold, although the underlying values other than the selected point were not numerically reported.

#### 4.1.2 Architectural ablation

Table 2 collects every ablation value reported in the source study. The reporting is incomplete: the source refers to an ablation table that is not included in the report and gives only selected values in its text, so each variant has between one and four reported numbers and the variants share few common columns. The evidence therefore supports qualitative statements about the direction of each change rather than a complete attribution of performance to individual components.

**Table 2.** Study 1 architectural ablation (BraTS 2020, 2D). Dice values exactly as reported in the source study. The source did not provide a complete ablation table; cells marked NR were not reported and have not been estimated. All values are from single training runs.

| Variant | Components |  | Dice |  |  |  |
| --- | --- | --- | --- | --- | --- | --- |
|  | SIB blocks<sup>a</sup> | Attention gates | Whole/overall<sup>b</sup> | NEC | ED | ET |
| U-Net (baseline) | – | – | 0.6772 | NR | 0.7080 | NR |
| SIB-UNet | ✓ | – | 0.7580 | NR | 0.8076 | NR |
| Attention U-Net | – | ✓ | NR | NR | NR | 0.8265 |
| SIBA-UNet (full) | ✓ | ✓ | **0.8601<sup>c</sup>** | 0.8795 | 0.8501 | 0.9125 |

<sup>a</sup> Separable inverted-bottleneck encoder and decoder blocks (Eqs. (3) and (4)).
<sup>b</sup> For the two baselines, the source reports this value as whole-tumor Dice. For SIBA-UNet, it reports an “overall” Dice, which it lists under the whole-tumor column of its literature table; the aggregation was not documented in enough detail to reconstruct it. It is not the arithmetic mean of the NEC, ED and ET values. The column is therefore labelled as reported and should not be read as a strictly like-for-like comparison.
<sup>c</sup> Validation Dice at the validation-selected threshold of 0.51 (Fig. 4b). The source does not state the partition for the other entries; given its statement that all results were obtained on a custom validation split, they are most plausibly validation values as well.

Three observations can be made from the reported values. First, replacing standard convolutions with separable inverted-bottleneck blocks (SIB-UNet) was associated with higher whole-tumor Dice than the baseline U-Net (0.7580 vs. 0.6772). The largest reported per-region change was for edema (0.7080 to 0.8076, +0.0996), a spatially diffuse region. Second, adding attention gates to an otherwise standard U-Net (Attention U-Net) was reported to improve the enhancing tumor, which reached an ET Dice of 0.8265. The source discusses less benefit from attention alone for edema but does not provide a numerical value. Third, the full SIBA-UNet, combining both modifications, reported the highest value for every region with a reported comparison: NEC 0.8795, ED 0.8501, ET 0.9125, and an overall Dice of 0.8601.

Two caveats apply. The “overall” value for SIBA-UNet and the whole-tumor values for the baselines are reported in the same column of the source’s own comparison, but the source does not document that they were computed identically (Table 2, note b). In addition, the reported ET Dice of 0.9125 exceeds the NEC and ED values. In BraTS, the enhancing tumor is commonly the most difficult region to segment, so this ordering is atypical. Without a documented Dice-aggregation protocol (for example, the handling of slices in which ET is absent), we cannot determine whether it reflects model behaviour or the evaluation procedure. We therefore report it without further interpretation.

#### 4.1.3 Qualitative segmentation

Representative outputs of the four Study 1 variants for a single axial slice are shown in Fig. 5, together with the target (reference) overlay. Consistent with the ablation, the variants differ most visibly in spatial specificity. The baseline U-Net produced diffuse, low-intensity probability maps in all three channels. The SIB-UNet edema map was more compact and ring-shaped than the baseline’s. Both attention-equipped variants (Attention U-Net and SIBA-UNet) produced compact, annular enhancing-tumor maps and a small, localised NEC response. Differences between Attention U-Net and SIBA-UNet on this slice are subtle. This is a single representative example: case identity and partition membership were not recorded in the source, and the figure illustrates the reported trends without demonstrating them at the population level.

**Figure 5.**
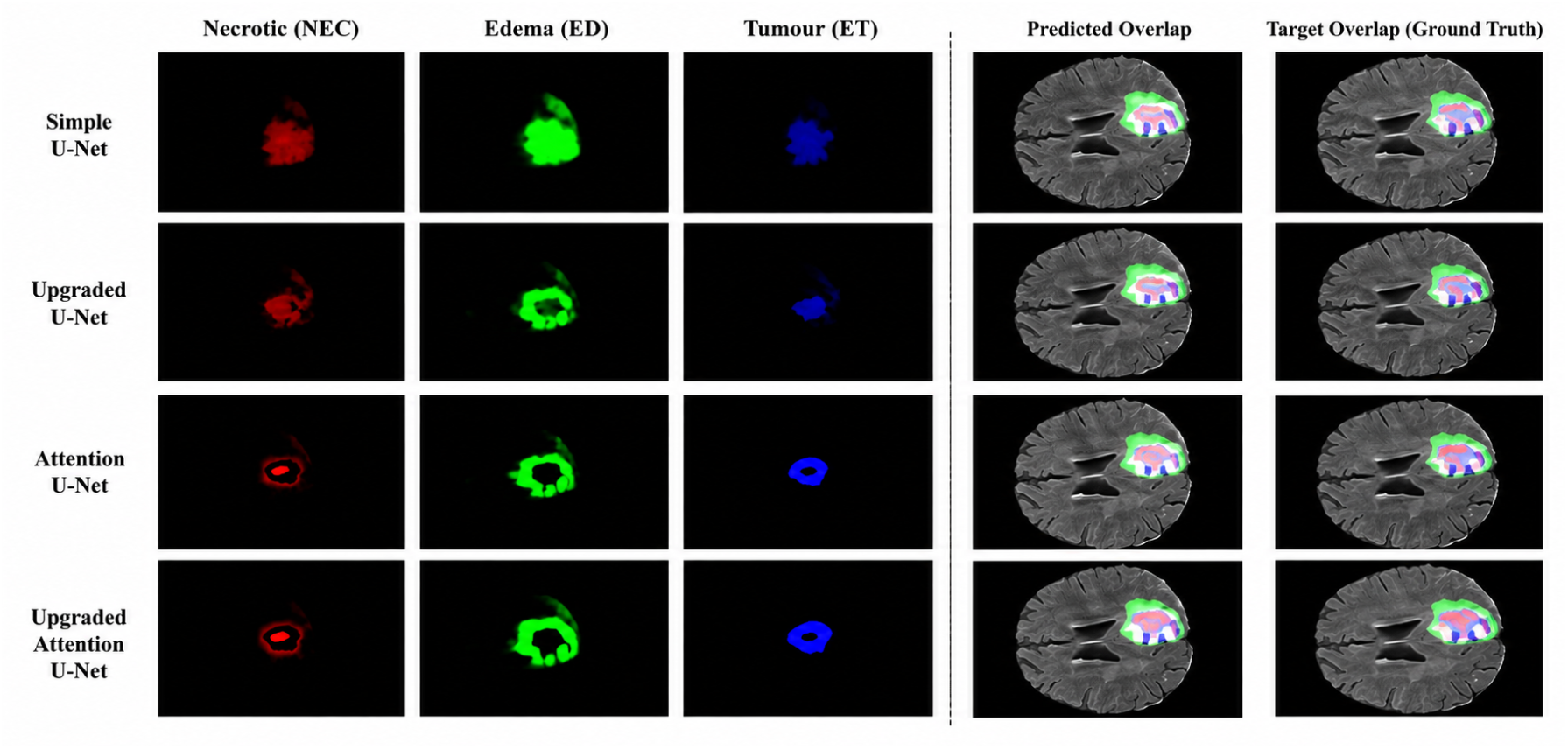
Study 1 qualitative comparison of architectural variants (representative BraTS 2020 axial slice). Rows: baseline U-Net (“Simple U-Net”), SIB-UNet (separable inverted-bottleneck blocks without attention gates), Attention U-Net, and SIBA-UNet. Columns: per-channel sigmoid probability maps for NEC, ED and ET (red, green and blue), followed by the predicted overlay and the target (reference) overlay on the corresponding MRI slice. The panels are re-composed from the original Study 1 outputs. Row labels in the image use the working names of the original study: “Simple U-Net” = baseline U-Net, “Upgraded U-Net” = SIB-UNet, “Attention U-Net” = Attention U-Net, “Upgraded Attention U-Net” = SIBA-UNet. Case identity and partition were not recorded.

### 4.2 Study 2: Physiologically informed 3D segmentation

#### 4.2.1 Input-representation ablation

Table 3 reports Dice and HD95 for the four input configurations, all trained and evaluated with the identical 3D U-Net and the identical case split. The 4-channel baseline achieved Dice scores of 0.8801 (WT), 0.8527 (TC) and 0.8273 (ET), for a mean Dice of 0.8534.

**Table 3.** Study 2 input-representation ablation with a fixed 3D U-Net (BraTS-GLI 2023). Held-out evaluation results exactly as reported in the source study; the architecture, preprocessing, split (1,001/125/125 cases), optimiser and checkpoint rule were identical across rows. Dice: higher is better; HD95: lower is better. Best value per column in bold. Single training run per configuration.

| Configuration<sup>a</sup> | Dice |  |  |  | HD95<sup>b</sup> |  |  |  |
| --- | --- | --- | --- | --- | --- | --- | --- | --- |
|  | WT | TC | ET | Mean | WT | TC | ET | Mean |
| 4-channel baseline | 0.8801 | 0.8527 | 0.8273 | 0.8534 | 8.2612 | 8.6076 | 7.8235 | 8.2308 |
| 6-channel Set A | 0.8807 | 0.8578 | 0.8231 | 0.8539 | 9.3215 | 8.5066 | 7.9445 | 8.5909 |
| 6-channel Set B | 0.8861 | 0.8593 | 0.8259 | 0.8571 | 8.4367 | 8.6128 | 7.6674 | 8.2390 |
| 8-channel (CMSR-3D) | **0.9101** | **0.8601** | **0.8396** | **0.8699** | **6.8801** | **6.6241** | **4.4536** | **5.9859** |
| $\Delta$ 8-channel – baseline<sup>c</sup> | +0.0300 | +0.0074 | +0.0123 | +0.0165 | –1.3811 | –1.9835 | –3.3699 | –2.2449 |

<sup>a</sup> Inputs: baseline, T1, T1CE, T2, FLAIR; Set A, baseline + T1–T1CE + T2–FLAIR; Set B, baseline + FLAIR–T1CE + T2–T1CE; 8-channel, baseline + all four difference maps.
<sup>b</sup> HD95 unit not specified in the source (voxels in the $128^3$ grid or millimetres).
<sup>c</sup> Differences computed by the present authors from the reported values; means are the arithmetic means of the three regions and reproduce the reported means exactly.

Adding either pair of difference maps alone produced small and mixed changes. Set A (T1*−*T1CE, T2*−*FLAIR) left mean Dice essentially unchanged (0.8539, +0.0005): it slightly increased WT and TC Dice, decreased ET Dice (0.8231), and increased WT HD95 from 8.2612 to 9.3215, which raised mean HD95 to 8.5909. Set B (FLAIR*−*T1CE, T2*−*T1CE) raised mean Dice to 0.8571 (+0.0037), mainly through WT (0.8861) and TC (0.8593); ET Dice (0.8259) remained slightly below the baseline, and mean HD95 (8.2390) was similar to the baseline (8.2308). Neither two-map set improved ET Dice.

The 8-channel configuration, which supplies all four difference maps (the full CMSR-3D representation), gave the highest reported mean Dice among the evaluated configurations (0.8699, +0.0165 over the baseline) and the highest Dice for each region: WT 0.9101, TC 0.8601 and ET 0.8396. It also gave the lowest HD95 for every region and the lowest mean HD95 (5.9859 vs. 8.2308). The largest boundary change was for ET, whose HD95 decreased from 7.8235 to 4.4536. Thus the full representation was the only configuration that improved ET on both metrics, and the only one that reduced HD95 substantially in all three regions.

The two 6-channel sets changed mean Dice by +0.0005 and +0.0037, whereas the 8-channel configuration changed mean Dice by +0.0165. The improvement observed with all four maps was therefore larger than the sum of the changes observed for the two partial configurations. Under the fixed architecture, this pattern is consistent with a possible joint benefit of the complete representation, although interaction effects were not formally tested. Each configuration was trained once, however, and run-to-run variability was not measured. The differences between the baseline and the two 6-channel sets are small enough that they may lie within training noise, and we do not interpret them further. The 8-channel differences are larger and consistent across all six region–metric combinations, but a single run cannot establish their reproducibility.

#### 4.2.2 Cross-modal subtraction feature analysis

Fig. 6 visualises the four difference maps for a representative BraTS-GLI 2023 case, with the reference subregion contours overlaid in axial, sagittal and coronal views. The colour bars indicate that intensities in this figure were rescaled to [0, 1] per modality for display, so the colour scales differ from the z-scored values seen by the network. In each map, red indicates that the first modality is relatively brighter and blue that the second is relatively brighter.

**Figure 6.**
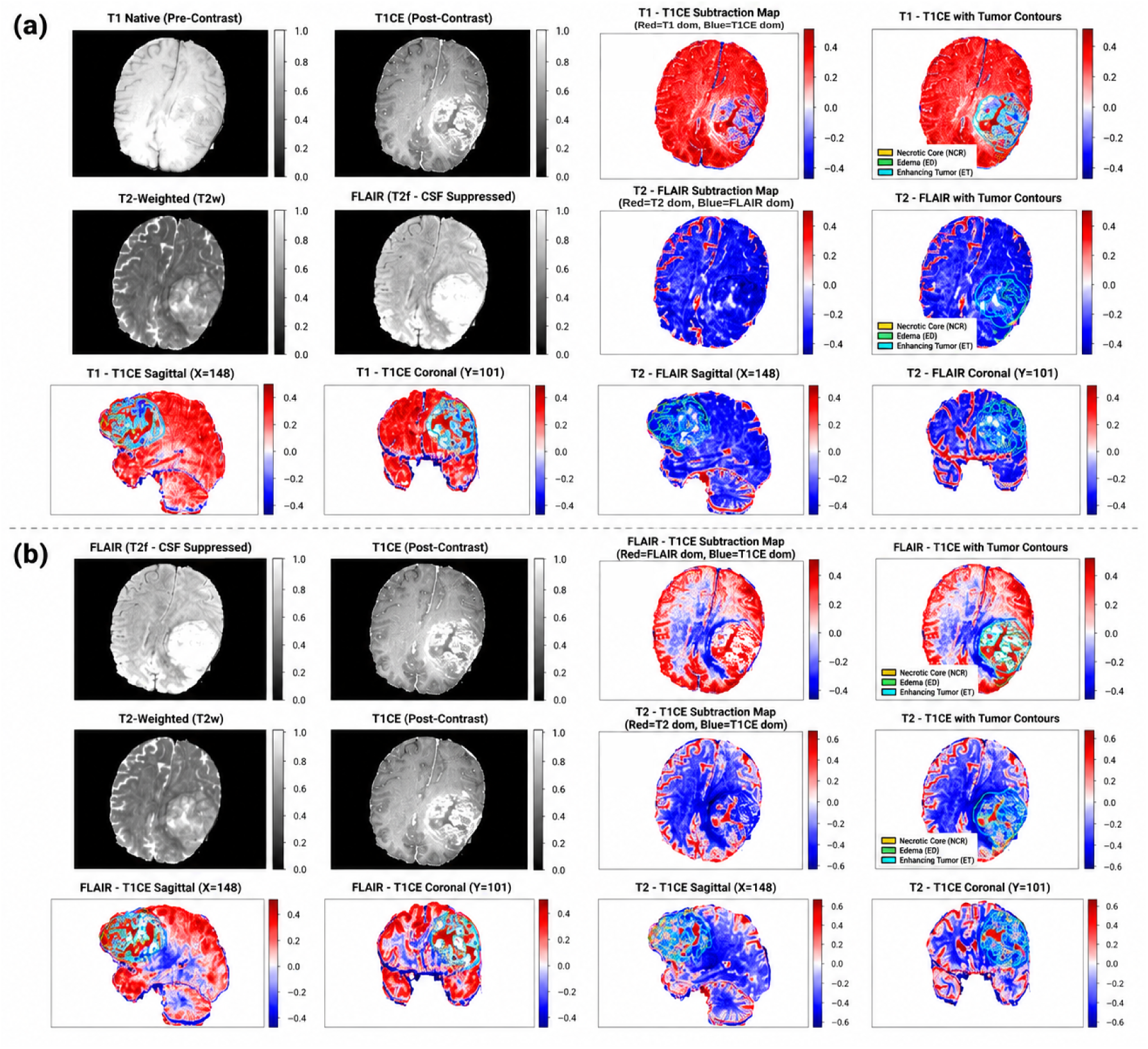
Cross-modal subtraction maps for a representative BraTS-GLI 2023 case. (a) Top two rows: T1 and T1CE with their difference map T1*−*T1CE, and T2 and FLAIR with their difference map T2*−*FLAIR, each map shown without and with reference contours (NCR, yellow; ED, green; ET, cyan). Bottom row: sagittal (*x* = 148) and coronal (*y* = 101) views of the two maps, with coordinates in the original image grid. (b) Same layout for FLAIR*−*T1CE and T2*−*T1CE. Colour bars on the difference maps: red, first modality dominant; blue, second modality dominant. Modality images are displayed on a 0–1 scale and differ from the z-scored network inputs. Case identity and partition were not recorded.

In this example, the T1*−*T1CE map (Fig. 6a, top) is predominantly red in normal-appearing brain and shows a concentrated blue pattern within the annotated tumor. The blue region lies within the annotated tumor contours, consistent with relative T1CE hyperintensity where contrast accumulates [14]. The T2*−*FLAIR map is predominantly blue in parenchyma and red in cerebrospinal-fluid spaces and sulci, as expected from FLAIR fluid suppression. Within the tumor region its contrast is weaker and more heterogeneous. The two T1CE-referenced maps (Fig. 6b) show mixed red and blue patterns inside the contours that differ between the core and the surrounding edema. The sagittal and coronal views show that these patterns are spatially coherent across planes rather than confined to a single slice.

These observations describe the maps of one representative case. The maps visualise intensity differences that spatially coincide with annotated subregions in this example; they are not validated physiological or biological measurements, and their behaviour across the cohort was not quantified.

#### 4.2.3 Boundary agreement (HD95)

HD95 changed more than Dice in relative terms. Relative to the baseline, the 8-channel configuration reduced mean HD95 by 2.2449 (27%), with reductions of 1.3811 (WT), 1.9835 (TC) and 3.3699 (ET, 43%). Dice, by contrast, rose by 0.0165 (1.9%). Because HD95 reflects the largest boundary deviations and is sensitive to small spurious components far from the tumor [27, 36], this pattern is consistent with a larger change in boundary-distance measurements than in bulk overlap, particularly for ET, although the specific error mechanisms were not analysed at the per-case level. Per-case distributions were not available to test this directly. Set A was the only configuration that worsened mean HD95, owing to its higher WT HD95. Because HD95 units were not reported, the values support comparisons between configurations within Study 2 but not comparisons with distances reported elsewhere.

#### 4.2.4 Qualitative, probability and boundary analysis

Fig. 7 presents one representative case from the 8-channel CMSR-3D model. The reference and predicted segmentations (Fig. 7a) agree closely in this slice: the necrotic core is enclosed by an enhancing rim, and a separate lateral edema crescent is reproduced. The visible discrepancies are small and lie mainly along the enhancing rim and the outer edema boundary. Panel (b) shows all eight input channels for the same slice, illustrating how the difference maps rearrange contrast relative to the source modalities; in particular, the enhancing rim appears as a dark band in the T1CE-referenced differences. The class probability maps (Fig. 7c) are spatially sharp: near-zero background probability across the lesion, a compact high-probability NCR region, an annular ET region, and high edema probability concentrated in the lateral crescent and along the rim. Intermediate probabilities occur mainly along class interfaces, and these interface regions are where the contour comparison (Fig. 7d) shows the residual differences between reference and predicted boundaries.

**Figure 7.**
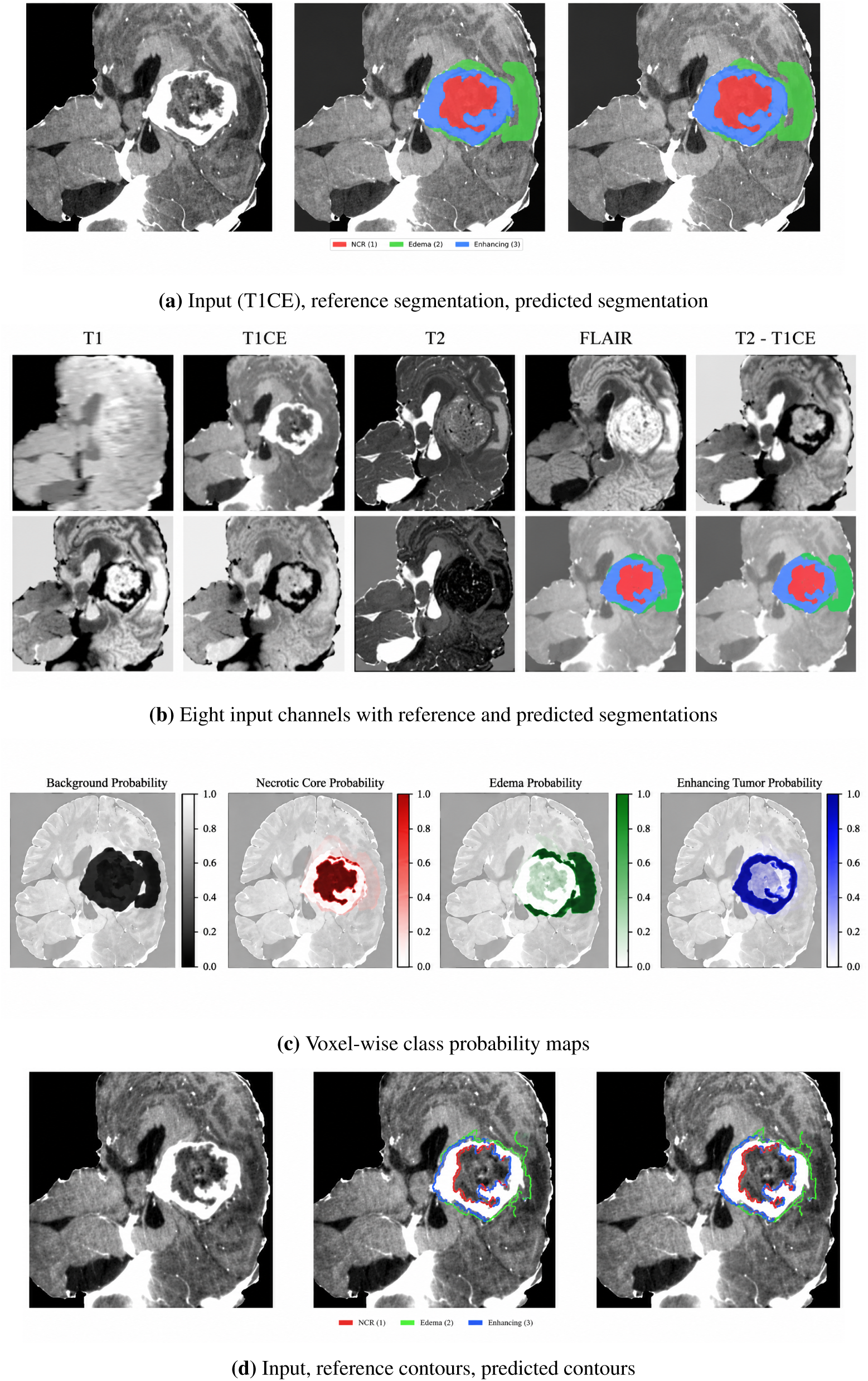
Representative Study 2 output of the 8-channel CMSR-3D model (BraTS-GLI 2023). (a) T1CE slice, reference segmentation and prediction. Left to right: input, reference segmentation, and predicted segmentation. (b) Top row: T1, T1CE, T2, FLAIR, T2*−*T1CE. Bottom row: FLAIR*−*T1CE, T1*−*T1CE, T2*−*FLAIR, reference segmentation, predicted segmentation. Bottom-row panels carry no labels in the image and are listed here from left to right. (c) Softmax probability maps for background, NCR, edema and enhancing tumor; each colour bar runs from 0 (no probability) to 1. (d) Reference and predicted subregion boundaries on the T1CE slice, in the same layout as (a). Colours: red, NCR; green, ED; blue, ET. Panels (a), (b) and (d) show the same slice; panel (c) is reproduced as displayed in the source figure. All panels are taken from the original output figures of Study 2. Case identity and partition were not recorded.

As with the other qualitative figures, this is a single example. It illustrates the model’s behaviour on one case and does not demonstrate robustness or performance across the cohort.

## 5 Cross-Study Analysis

### 5.1 Architecture-level versus representation-level design

A segmentation network can be written as *Ŷ* = *f_θ_* (*g*(*X*)), where *X* is the set of acquired MRI volumes, *g* maps them to the tensor presented to the network, and *f_θ_* is the network itself. The two studies intervene on different terms of this composition.

*Study 1 changes f_θ_.* The input mapping *g* is the conventional one: four normalised modalities stacked as channels of a 2D slice. The ablation variants differ only in how features are computed (standard versus separable inverted-bottleneck blocks) and how skip information is routed (direct versus attention-gated). The design question is how to spend a limited parameter and computation budget so that the network extracts and transmits the most useful features from a fixed input.

*Study 2 changes g.* The function class of *f_θ_* is fixed: the same 3D U-Net, apart from the width of its first convolution. The configurations differ only in whether, and which, explicit cross-modal contrasts are precomputed and supplied. The design question is whether making particular inter-sequence relationships explicit helps a fixed network learn them.

These are not two answers to the same question. The subtraction maps do not introduce new acquired measurements; rather, they provide explicitly constructed relationships between existing modalities as an input-space inductive bias. Their potential value is as an inductive bias: they present a relationship directly instead of requiring the network to discover it from limited data. Conversely, architectural changes cannot recover information that the input does not contain. The two levers therefore act on different parts of the learning problem and can, in principle, be combined.

### 5.2 What is and is not directly comparable

The studies share a goal—voxel or pixel-wise delineation of glioma subregions using the same four MRI modality types (T1, T1CE, T2 and FLAIR) and both separated training and evaluation data above the slice level. Beyond that, they differ in almost every element that determines a reported score:

- **Data.** BraTS 2020 versus BraTS-GLI 2023: different public collections, with different case counts, partitions and label conventions.
- **Dimensionality and context.** Study 1 sees a 128 *×* 128 slice with no through-plane context and a fixed slice window; Study 2 sees a 128^3^ volume with full 3D context but a fixed crop that may truncate peripheral tissue.
- **Output formulation.** Study 1 predicts three independent sigmoid channels, so a pixel can carry several labels. Study 2 predicts four mutually exclusive softmax classes, from which nested regions are derived.
- **Evaluation targets.** Study 1 reports NEC, ED, ET and an “overall” Dice whose aggregation is not documented. Study 2 reports the nested regions WT, TC and ET, which are unions of classes. Even identically named targets (ET) are computed on different units (2D slices versus 3D volumes) under different aggregation rules.
- **Operating point and partition.** Study 1 reports validation results at a threshold selected on the same validation data; Study 2 reports held-out results with a checkpoint selected on validation data.

Consequently, no statement of the form “the 3D model outperformed the 2D model” or vice versa is supported, and none is made. The reported Study 1 overall Dice and the Study 2 8-channel TC Dice happen to share the same value; this is a numerical coincidence between unrelated quantities. What can be compared across the studies is the *structure of the evidence*: which variable was manipulated, what was held fixed, and how completely the outcome was documented.

### 5.3 Complementary design lessons

Read side by side, the two studies suggest three lessons that neither study yields alone.

#### Diffuse and compact subregions respond to different levers

In Study 1, the largest reported architectural gain from the separable inverted-bottleneck blocks was for edema, a spatially extensive region, whereas attention gates were reported to help mainly the compact enhancing tumor. In Study 2, the enhancing tumor was the region on which partial input changes were least helpful: neither two-map set improved ET Dice. Only the complete representation improved ET, which also showed the largest HD95 reduction. In both settings, the reported improvement for the enhancing tumor—a thin, enhancement-defined structure—was tied to a specific intervention: attention-based localisation in one case, and the complete set of enhancement-related and T1CE-referenced contrasts in the other. This parallel is qualitative and rests on incompletely reported Study 1 values, but it points to the enhancing tumor as the subregion most sensitive to design choices of either kind.

#### The strength of evidence differs between the levers

Study 2 varied one factor under a fixed architecture, fixed split and fixed training protocol, and reported every cell of its ablation. Within the limits of single runs, it isolates the contribution of the input representation. Study 1 changed two architectural components and reported only part of its factorial comparison, so the individual contributions of the separable blocks and of the attention gates are documented incompletely. Within the scope of the reported experiments, Study 2 provides more direct evidence about its own design variable because the input representation was varied while the architecture, split and training protocol were held fixed. This difference reflects reporting completeness and experimental control, not the relative importance of the two levers.

#### The levers are composable but were not composed

Nothing in either design prevents applying an efficiency-oriented, attention-gated architecture to a subtraction-augmented input, in 2D or 3D. Such a combination would require new experiments with both factors varied under one protocol, and its outcome cannot be inferred from the present results. The value of separating the levers is that it makes clear which variable a given experiment can speak to.

## 6 Discussion

This manuscript set out to examine two completed brain tumor segmentation studies through a single lens: which part of the segmentation pipeline was changed, and what the evidence shows about that change. The central observation is that performance can be pursued through at least two distinct design levers—the network architecture and the input representation—and that they pose different questions, require different experimental controls, and yield different kinds of evidence. Much of the segmentation literature reports improvements that combine architectural, representational and training changes in a single comparison with prior methods. Separating the levers is a simple step, but it clarifies what each result can and cannot support.

For the representation lever, Study 2 addresses a controlled question: whether explicit cross-modal subtraction maps improve segmentation when the underlying 3D U-Net, data split and training protocol are held fixed. The 8-channel configuration achieved the highest reported mean Dice (0.8699) and the lowest reported mean HD95 (5.9859) among the four evaluated input configurations, whereas the two 6-channel configurations produced only small and mixed changes. The mean-Dice improvement relative to the 4-channel baseline was +0.0005 for Set A, +0.0037 for Set B, and +0.0165 for the full 8-channel representation. The full representation also produced the highest Dice and lowest HD95 for each of the three evaluation regions. These findings are consistent with a benefit from providing the complete set of explicitly constructed cross-modal relationships to the fixed network, but the experiments do not establish a specific mechanism for that benefit. Because each configuration was trained only once, run-to-run variability and statistical significance could not be assessed. The term *physiologically motivated* refers to the radiological contrast relationships that motivated the choice of the subtraction maps [14, 15]; the study does not establish that the learned representations are physiologically meaningful measurements.

For the architecture lever, Study 1 shows that an efficiency-oriented redesign of a 2D U-Net—separable inverted-bottleneck blocks [11, 12] together with attention-gated skip fusion [9, 10]—was associated with higher reported Dice than the baseline and single-component variants for the regions on which they can be compared. The reported values are consistent with the components having complementary roles: the block redesign was associated with the largest gain for diffuse edema, and attention with the enhancing tumor. However, the incomplete ablation means that this division of labour is suggested rather than demonstrated. The design is efficiency-oriented in construction, because depthwise separable convolutions reduce the cost of each convolution relative to a standard one at equal width [24]. Its actual parameter count, memory use and runtime were not measured relative to the baselines, so no quantitative efficiency claim is made. For 2D slice-based segmentation, such efficiency is the main practical argument for the approach, since slice-wise models trade volumetric context for lower memory requirements [26]; documenting it quantitatively is the most important missing piece of Study 1.

Taken together, the two studies illustrate how the choice of experimental design determines what can be learned. Study 2’s one-factor ablation under a fixed architecture provides more direct evidence about its own variable because the input representation was the only experimental factor varied. This is a statement about experimental control within a single run per configuration, not a formal causal-inference claim. It also says nothing about which lever is more important in general. Study 1 provides less complete evidence for component-level attribution because only part of its architectural ablation was reported. The comparison underlines a point made in recent methodological guidance for medical image analysis: controlled ablations, documented aggregation, and variance across runs or folds are what allow a reader to attribute an improvement to a design choice [27–29]. The reporting conventions adopted here (Section 3.5) follow from that guidance and from the CLAIM checklist [30, 31].

Several directions follow directly. The representation result invites a factorial study in which subtraction-augmented inputs are combined with different architectures—including efficiency-oriented and attention-gated designs, nnU-Net-style configurations [8] and transformer-based models [13]—under one protocol and with repeated runs. That study would show whether the input-level benefit persists when the architecture is stronger, a question the present evidence cannot answer. The individual difference maps could also be ablated singly, and normalised difference maps could be compared with raw differences. On the architecture side, a complete ablation with matched parameter budgets and measured cost would turn the efficiency-oriented design into a quantified efficiency result.

## 7 Limitations

The limitations below define the scope of what this manuscript establishes.

### Integration of two independent studies

The manuscript integrates two studies that differ in dataset release (BraTS 2020 and BraTS-GLI 2023), dimensionality (2D slices and 3D volumes), output formulation (multi-label sigmoid and mutually exclusive softmax), label conventions, data partitions and evaluation procedures. No model was trained jointly across the studies, no experiment combined the two design levers, and no numerical result is compared across the studies. The cross-study analysis concerns design logic and the structure of the evidence only. The two studies use different BraTS releases, with different datasets, partitions, preprocessing pipelines and evaluation protocols; consequently, their numerical results are not directly comparable.

### Study 1 reporting

The source report of Study 1 does not include a complete ablation table, so several variant–region combinations are unavailable (Table 2). It also does not specify the loss function, the channel widths and expansion factor, the kernel sizes, or the exact patient counts per partition. The aggregation underlying the “overall” Dice of 0.8601 could not be recovered. The reported results are validation results, with the decision threshold selected on the same validation partition, which makes the reported overall Dice an optimistically selected estimate. No held-out test result was reported. The 2D formulation uses a fixed 75-slice window without through-plane context. Parameter counts, computational cost and runtime were not measured, so the efficiency-oriented design is not supported by a quantitative efficiency benchmark.

### Study 2 reporting

The source of Study 2 documents a reserved test partition but does not state unambiguously whether its ablation table was computed on the validation or the test partition. The HD95 unit, the treatment of empty predictions in HD95, and the coordinate space in which metrics were computed were not reported. The split was described as subject-level, but BraTS-GLI identifiers include time-point suffixes, and the source does not state whether multiple time-points of the same patient were kept in the same partition. If they were not, longitudinal scans of one patient could appear in both training and evaluation data, which would bias results optimistically. The centre crop to 128^3^ may exclude peripheral tumor tissue in large or eccentric lesions. No data augmentation was used.

### Statistical evidence

Each configuration in both studies was trained once. No repeated runs, cross-validation folds, confidence intervals or significance tests are available, so the reproducibility of the reported differences is unknown [29]. This mainly affects the smaller differences, such as Study 2 Set A versus the baseline.

### Qualitative evidence

All qualitative figures show single representative examples for which case identity and partition membership were not recorded. They illustrate model behaviour and do not demonstrate robustness or generalisation. The subtraction-map interpretations describe intensity patterns in one case and are not validated physiological measurements.

### Figure provenance

Several result figures (Figs. 4 to 7) are re-rendered or re-composed from the original study outputs rather than reproduced directly, and the Study 1 loss and threshold curves are redrawn from the original plots; only the selected threshold point (0.51, Dice 0.8601) is numerically documented. Case identity and partition were not recorded for any qualitative example.

### External validity and reproducibility

Both studies used internal splits of a single public benchmark. No external, multi-institutional or prospective clinical validation was performed, and no claim of clinical utility is made. The original training implementations are no longer available, so the experiments cannot currently be re-executed from the authors’ code. The methodological descriptions in Section 3 are intended to be complete enough for independent reimplementation within the limits stated above.

## 8 Conclusion

This manuscript integrated two completed brain tumor segmentation studies around a single distinction: whether a study changes the network or the input. In the architecture-level study, SIBA-UNet—a 2D U-Net with separable inverted-bottleneck blocks and attention-guided skip fusion—reached an overall validation Dice of 0.8601 on BraTS 2020 at a validation-selected threshold, with reported gains over its single-component variants, although its ablation was only partially documented. In the representation-level study, CMSR-3D showed that, with a 3D U-Net held fixed, supplying all four cross-modal subtraction maps was associated with the highest Dice and the lowest HD95 in every region of BraTS-GLI 2023, whereas partial subsets were not.

The two levers act on different parts of the segmentation problem, can be studied separately, and in principle can be combined. Their results are not interchangeable, and they are not compared numerically here. The controlled, single-variable design of the representation study makes it the clearer evidence about its own variable. The main next steps are to confirm the representation effect with repeated runs and other architectures, and to complete the architectural ablation with measured computational cost.

## Declarations

### Ethics statement

This work used only publicly available, de-identified MRI data from the BraTS 2020 and BraTS 2023 adult glioma (BraTS-GLI) collections, released for research use by the challenge organisers [2–5]. No new data were collected from human participants, and the authors had no access to identifiable information. As the analyses were performed using openly available, de-identified secondary data, no additional participant recruitment or prospective human-subject procedures were undertaken.

### Data availability

The datasets analysed in this study are publicly available from the BraTS challenge organisers, subject to their data-use terms: the BraTS 2020 training data and the BraTS 2023 adult glioma (BraTS-GLI) training data, as described in the BraTS reference publications [2–5]. No new data were generated. All numerical results reported in this manuscript are contained in Tables 2 and 3. Lists of the case identifiers assigned to each partition are not available. The documented Study 2 partitioning procedure (sorting identifiers, shuffling with seed 42, and an 80:10:10 split) is provided in the Methods to facilitate replication.

### Author contributions

Latchan Chhetri led the conceptualisation, methodology, formal analysis, visualisation, and preparation of the manuscript, including the original draft and subsequent review and editing. Ayush Anand contributed to the conceptualisation, methodology, software, investigation, visualisation, and review and editing of the manuscript. Both authors reviewed and approved the final manuscript. Latchan Chhetri is the corresponding author.

## Acknowledgements

The authors sincerely thank Dr. Palash Ghosal for his teaching in biomedical image analysis, his valuable advice, and the insightful discussions that helped shape our understanding and interest in medical imaging.

## Use of generative AI

Generative AI was used to assist with the formatting of this manuscript. The authors reviewed and take responsibility for the final manuscript.

## Reporting guideline

This study was reported with reference to the Checklist for Artificial Intelligence in Medical Imaging (CLAIM 2024) [31].

## References

[1] David N. Louis, Arie Perry, Pieter Wesseling, Daniel J. Brat, Ian A. Cree, Dominique Figarella-Branger, et al. The 2021 WHO classification of tumors of the central nervous system: a summary. Neuro-Oncology, 23(8):1231–1251, 2021. doi: 10.1093/neuonc/noab106.

[2] Bjoern H. Menze, Andras Jakab, Stefan Bauer, Jayashree Kalpathy-Cramer, Keyvan Farahani, Justin Kirby, et al. The multimodal brain tumor image segmentation benchmark (BRATS). IEEE Transactions on Medical Imaging, 34(10):1993–2024, 2015. doi: 10.1109/TMI.2014.2377694.

[3] Spyridon Bakas, Hamed Akbari, Aristeidis Sotiras, Michel Bilello, Martin Rozycki, Justin S. Kirby, John B. Freymann, Keyvan Farahani, and Christos Davatzikos. Advancing The Cancer Genome Atlas glioma MRI collections with expert segmentation labels and radiomic features. Scientific Data, 4: 170117, 2017. doi: 10.1038/sdata.2017.117.

[4] Spyridon Bakas, Mauricio Reyes, Andras Jakab, Stefan Bauer, Markus Rempfler, Alessandro Crimi, et al. Identifying the best machine learning algorithms for brain tumor segmentation, progression assessment, and overall survival prediction in the BRATS challenge. arXiv:1811.02629, 2018.

[5] Ujjwal Baid, Satyam Ghodasara, Suyash Mohan, Michel Bilello, Evan Calabrese, Errol Colak, et al. The RSNA-ASNR-MICCAI BraTS 2021 benchmark on brain tumor segmentation and radiogenomic classification. arXiv:2107.02314, 2021.

[6] Olaf Ronneberger, Philipp Fischer, and Thomas Brox. U-Net: Convolutional networks for biomedical image segmentation. In Medical Image Computing and Computer-Assisted Intervention – MICCAI 2015, volume 9351 of Lecture Notes in Computer Science, pages 234–241. Springer, 2015. doi: 10.1007/978-3-319-24574-4_28.

[7] Özgün Çiçek, Ahmed Abdulkadir, Soeren S. Lienkamp, Thomas Brox, and Olaf Ronneberger. 3D U-Net: Learning dense volumetric segmentation from sparse annotation. In Medical Image Computing and Computer-Assisted Intervention – MICCAI 2016, volume 9901 of Lecture Notes in Computer Science, pages 424–432. Springer, 2016. doi: 10.1007/978-3-319-46723-8_49.

[8] Fabian Isensee, Paul F. Jaeger, Simon A. A. Kohl, Jens Petersen, and Klaus H. Maier-Hein. nnU-Net: a self-configuring method for deep learning-based biomedical image segmentation. Nature Methods, 18 (2):203–211, 2021. doi: 10.1038/s41592-020-01008-z.

[9] Ozan Oktay, Jo Schlemper, Loic Le Folgoc, Matthew Lee, Mattias Heinrich, Kazunari Misawa, et al. Attention U-Net: Learning where to look for the pancreas. arXiv:1804.03999, 2018.

[10] Jo Schlemper, Ozan Oktay, Michiel Schaap, Mattias Heinrich, Bernhard Kainz, Ben Glocker, and Daniel Rueckert. Attention gated networks: Learning to leverage salient regions in medical images. Medical Image Analysis, 53:197–207, 2019. doi: 10.1016/j.media.2019.01.012.

[11] François Chollet. Xception: Deep learning with depthwise separable convolutions. In Proceedings of the IEEE Conference on Computer Vision and Pattern Recognition (CVPR), pages 1800–1807, 2017. doi: 10.1109/CVPR.2017.195.

[12] Mark Sandler, Andrew Howard, Menglong Zhu, Andrey Zhmoginov, and Liang-Chieh Chen. MobileNetV2: Inverted residuals and linear bottlenecks. In Proceedings of the IEEE/CVF Conference on Computer Vision and Pattern Recognition (CVPR), pages 4510–4520, 2018. doi: 10.1109/CVPR.2018.00474.

[13] Ali Hatamizadeh, Vishwesh Nath, Yucheng Tang, Dong Yang, Holger R. Roth, and Daguang Xu. Swin UNETR: Swin transformers for semantic segmentation of brain tumors in MRI images. In Brainlesion: Glioma, Multiple Sclerosis, Stroke and Traumatic Brain Injuries (BrainLes 2021), volume 12962 of Lecture Notes in Computer Science, pages 272–284. Springer, 2022. doi: 10.1007/978-3-031-08999-2_22.

[14] Benjamin M. Ellingson, Hyun J. Kim, Davis C. Woodworth, Whitney B. Pope, et al. Recurrent glioblastoma treated with bevacizumab: contrast-enhanced T1-weighted subtraction maps improve tumor delineation and aid prediction of survival in a multicenter clinical trial. Radiology, 271(1): 200–210, 2014. doi: 10.1148/radiol.13131305.

[15] Sohil H. Patel, Laila M. Poisson, Daniel J. Brat, et al. T2–FLAIR mismatch, an imaging biomarker for IDH and 1p/19q status in lower-grade gliomas: A TCGA/TCIA project. Clinical Cancer Research, 23 (20):6078–6085, 2017. doi: 10.1158/1078-0432.CCR-17-0560.

[16] Lena Maier-Hein, Matthias Eisenmann, Annika Reinke, Sinan Onogur, Marko Stankovic, Patrick Scholz, et al. Why rankings of biomedical image analysis competitions should be interpreted with care. Nature Communications, 9:5217, 2018. doi: 10.1038/s41467-018-07619-7.

[17] Fausto Milletari, Nassir Navab, and Seyed-Ahmad Ahmadi. V-Net: Fully convolutional neural networks for volumetric medical image segmentation. In 2016 Fourth International Conference on 3D Vision (3DV), pages 565–571. IEEE, 2016. doi: 10.1109/3DV.2016.79.

[18] Zongwei Zhou, Md Mahfuzur Rahman Siddiquee, Nima Tajbakhsh, and Jianming Liang. UNet++: A nested U-Net architecture for medical image segmentation. In Deep Learning in Medical Image Analysis and Multimodal Learning for Clinical Decision Support (DLMIA 2018), volume 11045 of Lecture Notes in Computer Science, pages 3–11. Springer, 2018. doi: 10.1007/978-3-030-00889-5_1.

[19] Mohammad Havaei, Axel Davy, David Warde-Farley, Antoine Biard, Aaron Courville, Yoshua Bengio, Chris Pal, Pierre-Marc Jodoin, and Hugo Larochelle. Brain tumor segmentation with deep neural networks. Medical Image Analysis, 35:18–31, 2017. doi: 10.1016/j.media.2016.05.004.

[20] Konstantinos Kamnitsas, Christian Ledig, Virginia F. J. Newcombe, Joanna P. Simpson, Andrew D. Kane, David K. Menon, Daniel Rueckert, and Ben Glocker. Efficient multi-scale 3D CNN with fully connected CRF for accurate brain lesion segmentation. Medical Image Analysis, 36:61–78, 2017. doi: 10.1016/j.media.2016.10.004.

[21] Andriy Myronenko. 3D MRI brain tumor segmentation using autoencoder regularization. In Brainlesion: Glioma, Multiple Sclerosis, Stroke and Traumatic Brain Injuries (BrainLes 2018), volume 11384 of Lecture Notes in Computer Science, pages 311–320. Springer, 2019. doi: 10.1007/978-3-030-11726-9_28.

[22] Chen-Yu Lee, Saining Xie, Patrick Gallagher, Zhengyou Zhang, and Zhuowen Tu. Deeply-supervised nets. In Proceedings of the Eighteenth International Conference on Artificial Intelligence and Statistics (AISTATS), volume 38 of Proceedings of Machine Learning Research, pages 562–570, 2015.

[23] Dmitry Ulyanov, Andrea Vedaldi, and Victor Lempitsky. Instance normalization: The missing ingredient for fast stylization. arXiv:1607.08022, 2016.

[24] Andrew G. Howard, Menglong Zhu, Bo Chen, Dmitry Kalenichenko, Weijun Wang, Tobias Weyand, Marco Andreetto, and Hartwig Adam. MobileNets: Efficient convolutional neural networks for mobile vision applications. arXiv:1704.04861, 2017.

[25] Jeya Maria Jose Valanarasu and Vishal M. Patel. UNeXt: MLP-based rapid medical image segmentation network. In Medical Image Computing and Computer Assisted Intervention – MICCAI 2022, volume 13435 of Lecture Notes in Computer Science, pages 23–33. Springer, 2022. doi: 10.1007/978-3-031-16443-9_3.

[26] Arman Avesta, Sajid Hossain, MingDe Lin, Mariam Aboian, Harlan M. Krumholz, and Sanjay Aneja. Comparing 3D, 2.5D, and 2D approaches to brain image auto-segmentation. Bioengineering, 10(2): 181, 2023. doi: 10.3390/bioengineering10020181.

[27] Annika Reinke, Minu D. Tizabi, Michael Baumgartner, Matthias Eisenmann, Doreen Heckmann-Nötzel, A. Emre Kavur, et al. Understanding metric-related pitfalls in image analysis validation. Nature Methods, 21(2):182–194, 2024. doi: 10.1038/s41592-023-02150-0.

[28] Lena Maier-Hein, Annika Reinke, Patrick Godau, Minu D. Tizabi, Florian Buettner, Evangelia Christodoulou, et al. Metrics reloaded: recommendations for image analysis validation. Nature Methods, 21(2):195–212, 2024. doi: 10.1038/s41592-023-02151-z.

[29] Gaël Varoquaux and Veronika Cheplygina. Machine learning for medical imaging: methodological failures and recommendations for the future. npj Digital Medicine, 5:48, 2022. doi: 10.1038/s41746-022-00592-y.

[30] John Mongan, Linda Moy, and Charles E. Kahn. Checklist for artificial intelligence in medical imaging (CLAIM): A guide for authors and reviewers. Radiology: Artificial Intelligence, 2(2):e200029, 2020. doi: 10.1148/ryai.2020200029.

[31] Ali S. Tejani, Michail E. Klontzas, Anthony A. Gatti, John T. Mongan, Linda Moy, Seong Ho Park, Charles E. Kahn, et al. Checklist for artificial intelligence in medical imaging (CLAIM): 2024 update. Radiology: Artificial Intelligence, 6(4):e240300, 2024. doi: 10.1148/ryai.240300.

[32] Diederik P. Kingma and Jimmy Ba. Adam: A method for stochastic optimization. In 3rd International Conference on Learning Representations (ICLR), 2015. doi: 10.48550/arXiv.1412.6980.

[33] Ilya Loshchilov and Frank Hutter. Decoupled weight decay regularization. In 7th International Conference on Learning Representations (ICLR), 2019. doi: 10.48550/arXiv.1711.05101.

[34] Ilya Loshchilov and Frank Hutter. SGDR: Stochastic gradient descent with warm restarts. In 5th International Conference on Learning Representations (ICLR), 2017. doi: 10.48550/arXiv.1608.03983.

[35] Lee R. Dice. Measures of the amount of ecologic association between species. Ecology, 26(3):297–302, 1945. doi: 10.2307/1932409.

[36] Abdel Aziz Taha and Allan Hanbury. Metrics for evaluating 3D medical image segmentation: analysis, selection, and tool. BMC Medical Imaging, 15:29, 2015. doi: 10.1186/s12880-015-0068-x.

